# Individual differences in experiential diversity shape event segmentation granularity

**DOI:** 10.1101/2022.07.07.499122

**Authors:** Carl J. Hodgetts, Samuel C. Berry, Mark Postans, Angharad N. Williams

## Abstract

Parsing experience into meaningful events or units, known as event segmentation, may be critical for structuring episodic memory, planning, and navigating the spatial and social world. However, little is known about what factors shape inter-individual differences in event segmentation. Here, we show that individuals with greater variation in their daily social and spatial lives (experiential diversity) displayed more fine-grained event segmentation during a movie-viewing task. Further analyses revealed that this relationship held after considering potential confounds, such as anxiety and loneliness, and was primarily driven by variation in social experiential diversity. These results support the view that event segmentation can occur proactively based on social and spatial environmental dynamics learned ‘in the wild’ and provide a potential cognitive pathway through which isolation impacts cognitive health.

## Introduction

While the world provides a continuous stream of perceptual input, we perceive and remember our daily lives as a series of discrete events with distinct beginnings and endings. This process, known as event segmentation, is thought to rely on the detection of event boundaries^1,2^. These boundaries can be estimated implicitly (e.g., by clustering fMRI activity into discrete states during continuous movie viewing)^3,4^ or measured directly by asking participants to press a button whenever they perceive one meaningful unit of activity to end and another to begin^5^. While the precise neurocognitive mechanisms for determining event boundaries are still debated^6–8^, there is growing evidence that event segmentation strongly influences several other aspects of cognition, including spatial navigation, long-term memory, and motor planning^9–11^.

Despite its potentially broad psychological relevance, there remains a limited understanding of how, and why, this ability differs across individuals. While studies often highlight high agreement in where people place event boundaries^12^, there is nonetheless evidence showing substantial variation in the number of events individuals perceive for the same stimulus^6,13,14^ – a finding that is consistent with known individual differences in the granularity of event memories^15^.

As event representations are shaped and updated through interactions with the external world ^2^, one potential source of inter-individual variation could be the degree of situational change experienced in one’s physical and/or social environment. Indeed, people differ significantly in how much variability they encounter in their daily lives—referred to as ‘experiential diversity’^16,17^.

Evidence linking event processing to experiential diversity predominantly stems from studies in nonhuman species. For example, research on environmental and social enrichment in rodents suggest that experiential diversity is not only critical for psychological wellbeing (i.e., reducing stress) but promotes spatial-event memory alongside structural plasticity in brain regions associated with event processing in humans, such as the hippocampus and prefrontal cortex^18–20^. In humans, markers of low social experiential diversity, such as social isolation, have been associated with memory impairment in older adults^21^. Studies on spatial diversity indicate that individuals raised in areas with high street network entropy (i.e., more complex, less ordered environments) exhibit better navigation skills in adulthood^22^ (see also ref. ^23^). Similarly, studies involving both younger and older adults have shown that behavioural interventions that promote exploration within novel, large-scale environments have also been shown to improve mnemonic discrimination in both younger and older adults, indicating enhanced fine-grained encoding of events^23,24^. Collectively, these findings imply that exposure to rich, varied experiences may enhance event-related cognition and induce structural changes in brain regions implicated in event segmentation^3,25^. However, it remains unknown whether real-world inter-individual differences in daily experiences are associated with the granularity of event segmentation itself.

Notably, the expected direction of the relationship between experiential diversity and event segmentation is not clear based on current theories. On the one hand, higher experiential diversity may increase the granularity of event segmentation because regular exposure to complex physical and social environments could enhance sensitivity to changes in fine-grained situational features that comprise events (e.g., social and environmental cues)^26^ and their resulting internal states (e.g., goals and motivations)^17,27^. Conversely, from the perspective of Event Segmentation Theory^1^, low levels of real-world experiential diversity may increase event segmentation granularity due to more imprecise predictions about the world. Specifically, the lack of exposure to highly variable experiences may increase the probability that minor shifts in situational features and/or internal states cross an event horizon. Understanding the relationship between experiential diversity and event segmentation, therefore, is not only relevant for understanding real-world psychological effects of low experiential diversity, such as those stemming from social isolation, loneliness, and poor physical or mental health, but is important in the theoretical understanding of event segmentation as a cognitive process.

To address this, we examined the association between real-world experiential diversity and event segmentation granularity within a sample of 157 young healthy adults. For each participant, we assessed both social and spatial experiential diversity over the preceding 30-day period, allowing us to examine the differential contributions of each^28^. Partial COVID-19 restrictions at the time of testing in the United Kingdom ^29^, which placed restrictions on the size of social gatherings, access to work and education, and travel, also provided a unique opportunity to study the effects of more restricted experiential diversity within a young, healthy adult sample. In these same participants, we applied an event segmentation task in which they watched a short film—an abridged version of Alfred Hitchcock’s *Bang! You’re Dead* (see Figure 1)—and marked boundaries in their experience using an on-screen button ^25,30^. The main measure was the number of perceived event boundaries reported by each participant (see ref. ^6,26,31,32^). We hypothesised that higher experiential diversity would lead to more fine-grained event segmentation.

**Figure 1.**
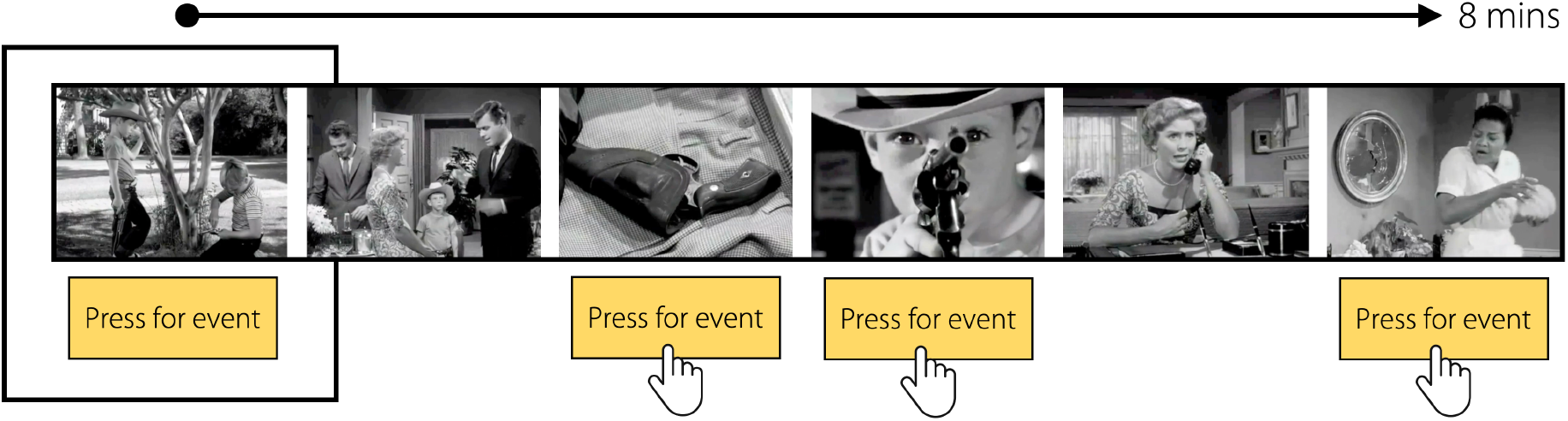
The event segmentation task. a) The event segmentation phase involved the presentation of an 8-minute movie: Alfred Hitchcock’s black-and-white television drama “Bang! You’re Dead”. The movie was shown above an on-screen button, which participants pressed whenever they judged one event (i.e., “a meaningful unit of time”) to end and another to begin.

## Methodology

### Participants

Participants were recruited from the participant recruitment platform Prolific (https://prolific.ac) and reimbursed for their participation at a rate of £8.49/hr. Participants were between 18-35 years of age, UK residents, fluent English speakers, and had no current or previous neurological or psychiatric conditions. Based on a priori power analysis, we aimed to collect a total of 153 complete datasets in a sample of young adult participants (which would provide 80% power to detect a one-tailed, small-to-moderate effect size of r = 0.2). A total of 177 participants engaged with the study on Prolific, and 157 provided complete data for all variables of interest (mean age = 26.4; SD = 5.4; range = 18-35). Participants’ gender was recorded using ‘female’ and ‘male’ options, with “Another gender not listed here” option where they could self-identify their gender. Ninety participants reported being female and 65 reported being male. One participant reported their gender as nonbinary, and one as trans. This study was approved by the Royal Holloway Department of Psychology Research Ethics Committee (ethical approval reference 2171).

### Event segmentation task

Participants from Prolific were directed to Qualtrics (www.qualtrics.com) to complete the experiment. The paradigm was optimised for computers, phones, and tablets. We used an event segmentation task that has been applied in recent functional MRI studies of event perception ^30^. Here, participants watched an abridged version of Alfred Hitchcock’s black-and-white television drama “Bang! You’re Dead”. This film was chosen as it is highly suspenseful, but also involves dynamic social interactions across multiple scenes and locations and is therefore ideal to examine the cognitive-perceptual consequences of social and spatial experiential diversity. This contrasts with many other studies of event segmentation that have mainly used videos depicting a single agent performing action sequences or daily activities ^5,33^. Importantly, no participant reported seeing this film before, meaning that perceived events were not driven by prior knowledge of the film itself. During the movie, participants were required to press an on-screen button (using their mouse or phone/tablet touchscreen) whenever they judged one event to end and another to begin (Figure 1). They were instructed that there were “no right answers in this task” and to just respond in a way that feels natural to them. The main measure of event segmentation was the total number of perceived event boundaries per participant (i.e., the total number of button presses during the movie)^6,26,31,32^.

### Experiential diversity

The experiential diversity measure was designed to capture the diversity of each participant’s ‘social’ and ‘spatial’ experiences over the preceding 30 days. The social experiential diversity score was comprised of two sub-scales. The first included several questions assessing the regularity and format of recent social interactions (see Supplementary Material, Appendix A). The second scale was a measure of social network size^34,35^, whereby participants provided the initials of every individual they had had meaningfully contacted over the last 30 days. This questionnaire is thought to probe the second layer of an individual’s social network (the ‘sympathy group’)^36^, and the 30-day limit is thought to maximise variation across individuals while also minimising the time and effort required to complete the questionnaire. The social experiential diversity score was a composite of these sub-scales and scores could range from 3-50 (mean = 15.9, SD = 8). The spatial experiential diversity questionnaire was designed to assess the complexity of each participant’s immediate environment, including the number of rooms they spend their time in on a typical day, access to private outdoor space, and the frequency in which they explore their local neighbourhood (see Supplementary Material, Appendix B). Scores on this measure could range from 0–16 (mean = 7.9, SD = 3, range in sample = 0-13). Scores on the social and spatial measures were combined into an overall experiential diversity score (mean = 23.8, SD = 9.3, range = 5-60).

### Other questionnaires

In addition to our main measures above, we also collected data on several covariates of interest. The first of these was The Campaign to End Loneliness Measurement Tool^37^, which is a 3-item tool to assess subjective feelings of loneliness. The three items on this tool are: *“1. I am content with my friendships and relationships”; “2. I have enough people I feel comfortable asking for help at any time”; “3. My relationships are as satisfying as I would want them to be”*. Participants could respond to each statement on a 5-point scale: ‘Strongly disagree’ (coded as 4 points), ‘Disagree’ (coded as 3 points), ‘Neutral’ (coded as 2 points), ‘Agree’ (coded as 1 point), and ‘Strongly agree’ (coded as 0 points). This coding scheme produces scores ranging from 0-12, where higher scores indicate higher levels of subjective loneliness. To control for general feelings of anxiety (see Results), participants also completed the trait component of the State-Trait Anxiety Inventory (STAI-T)^38^. The STAI-T contains 20 items assessing the frequency of anxiety generally felt by participants, and participants respond on a four-point scale from ‘Almost’ to ‘Almost Always’. Scores can range from 20-80 with high scores indicating high levels of general anxiety.

### Statistical analysis

All statistical analyses were conducted using R (version 4.0.2; R Core Team, 2019) in RStudio (version 1.4.1106; RStudio Team, 2021). Key correlational analyses (e.g., between experiential diversity and event segmentation granularity) were conducted using two-tailed bivariate correlations, with the contribution of potential covariates explored using partial correlations and multiple regression analyses (see Results). Partial correlations were carried out using the ‘ppcor’^39^ package in R. Comparisons between correlation coefficients were performed using Steiger Z-tests (Steiger, 1980) within the ‘cocor’ package in R^40^.

### Open Practices Statement

The design and analysis plan for the lab study were not preregistered. De-identified data, analysis code and task materials have been made available via the Open Science Framework and can be accessed freely at the following URL: https://osf.io/2q4wr/?view_only=e6f77d7ecca9427ca2847222bd67dfd9. Information on sample-size determination, data exclusion, and measure selection are reported in the respective Method sections above.

## Results

### Experiential diversity is strongly correlated with event segmentation

We predicted that experiential diversity would be positively associated with event segmentation granularity, or the number of event boundaries perceived during the movie stimulus. To test this, we first conducted bivariate Pearson’s correlations between total experiential diversity (i.e., social and spatial subscales combined) and event segmentation granularity (Figure 2). Consistent with our prediction, we observed a significant positive correlation between experiential diversity and event segmentation frequency (r (155) = 0.27, p = 0.0008).

**Figure 2.**
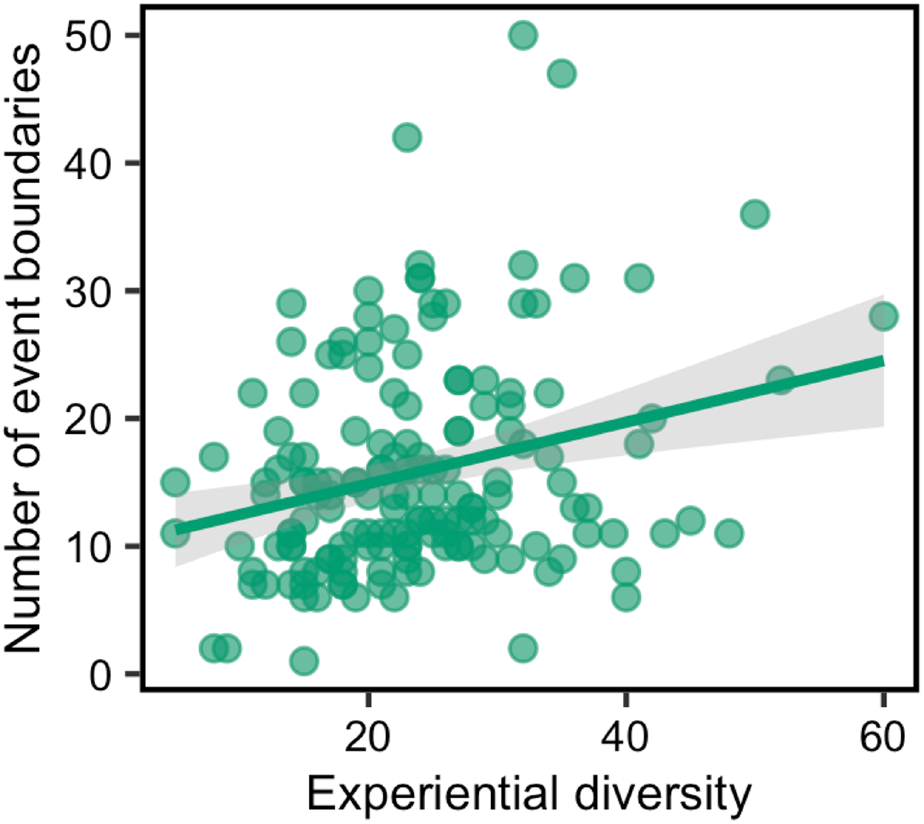
The relationship between total experiential diversity and event segmentation granularity. Each data point reflects an individual participant and there are 157 data points shown. The shaded area represents the 95% confidence interval on the best-fitting regression line.

### Controlling for anxiety and loneliness

One possible explanation for this finding is that low experiential diversity (e.g., smaller social networks and/or limited access to diverse spatial environments) is associated with higher levels of anxiety and loneliness. For example, over-general memory has been observed in mood disorders^41,42^, which may also be evident in how events are initially perceived (e.g., a bias towards *coarse*-grained event segmentation). To test this, we conducted a multiple regression with event segmentation as the outcome measure and experiential diversity, anxiety, and loneliness as the predictors (see Methods). Notably, the effect between experiential diversity and event segmentation held when controlling for both anxiety and loneliness (β = 0.25, p = 0.0008). Further, there were also no independent relationships between event segmentation granularity and either anxiety (p = 0.1) or loneliness (p = 0.19), suggesting that this effect is not driven by the subjective quality of individuals’ social interactions, or anxiety linked to isolation, but rather by the objective diversity of everyday social and spatial experiences.

### The differential contribution of ‘social’ and ‘spatial’ experiential diversity

Finally, we examined the relative contribution of social and spatial experiential diversity measures to the number of event boundaries segmented. We found that only social experiential diversity was significantly associated with event segmentation (*social*: r (155) = 0.25, p = 0.001; *spatial*: r (155) = 0.15, p = 0.06; see Figure 3), though these correlations did not differ significantly from one another (Z = 1.09, p = 0.28). An additional question is whether social or spatial components of experiential diversity make independent contributions to event segmentation. Indeed, when spatial experiential diversity is controlled for within in a multiple regression model, social experiential diversity still significantly accounts for the variation in event boundary perception (β = 0.24, p = 0.006), suggesting that social experiential diversity is the main contributor to event segmentation granularity in this study.

**Figure 3.**
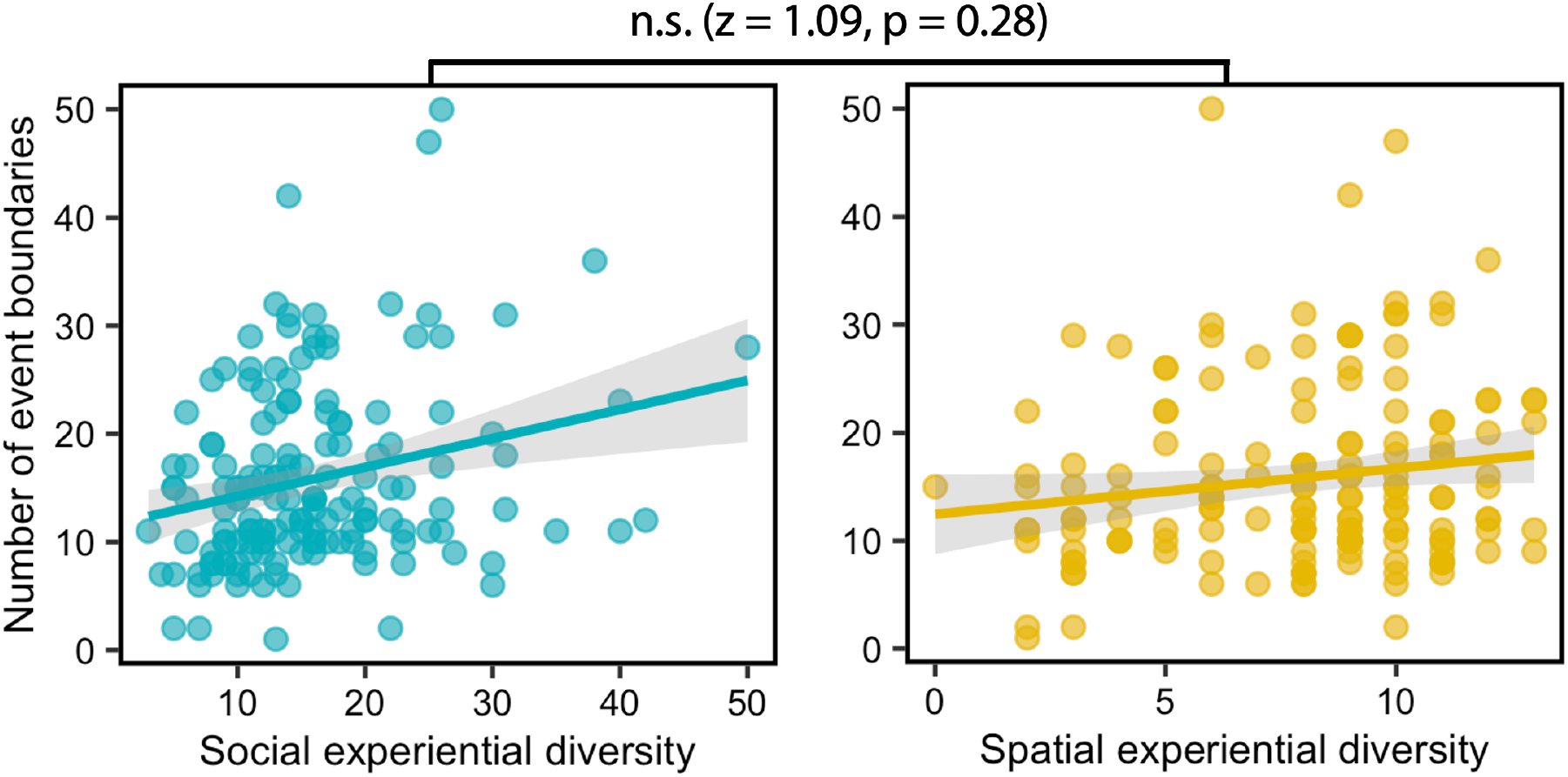
The relationship between social and spatial experiential diversity subscales and event segmentation granularity. Statistical comparisons between social and spatial conditions are reported above the plots (Steiger Z-test). Each data point reflects an individual participant and there are 157 data points shown. The shaded area represents the 95% confidence interval on the best-fitting regression line.

## Discussion

In this study, we investigated the association between inter-individual differences in real-world experiential diversity (e.g., the frequency and quality of social interactions, and the capacity to explore the immediate spatial environment) and inter-individual variation in event segmentation granularity. Our results yielded several key findings. First, we found a strong relationship between experiential diversity and the number of event boundaries perceived during a movie-viewing task, such that participants who had recent short-term exposure to a wider range of social and spatial experiences provided more fine-grained segmentations of events as they unfolded. Second, we found that this effect held when controlling for anxiety and loneliness, addressing the possibility that this effect reflects variation in mood/anxiety linked to isolation. Thirdly, when comparing social and spatial experiential diversity, we found that social experiences accounted for a greater proportion of variation in event perception compared to spatial experiences; however, a direct comparison between these correlations was not statistically significant.

The close relationship between the complexity/diversity of everyday experience and event segmentation granularity (as observed here) aligns with the view that event segmentation can be actively shaped by prior knowledge and expectations about environmental dynamics, rather than reflecting solely prediction error (see e.g., ref. ^8^ for recent empirical demonstration in the context of temporal order memory). From the perspective of Event Segmentation Theory, for example, it might be predicted that *low* experiential diversity (i.e., a state in which there are fewer major shifts in an individuals’ everyday environment) would lower the threshold for a prediction error spike to generate a new event model^1,43^, leading to overall greater sensitivity to event boundaries. We propose, however, that rather than reacting to prediction errors, event segmentation in individuals with high experiential diversity may operate more proactively, unitising experiences and internal states^17,27^ based on a more fine-grained understanding of the environment’s temporal dynamics and structure^7,8^ – as developed through experiential diversity.

This also aligns, in part, with research on event perception and expertise. For example, basketball experts generate more event boundaries compared to novices when watching basketball games, particularly when attending to fine-grained events^26^ (but see ref. ^44^). Thus, it may be that participants’ boundaries aligned at the coarse-level in the current study (i.e., for highly salient scene changes) but that individuals with higher experiential diversity had increased sensitivity to finer-scale, sub-events linked to changes in spatial and/or social features of events^9^. Such an interpretation could also be extended to internal states during movie viewing, such as emotional states, goals, and motivation^2,45^. Indeed, recent work has observed that individuals who have recently experienced more varied social contexts display more emotional granularity^17,27^. As such, it is possible that these effects reflect a combined contribution of varied sensitivity to both external cues and internal states (with the latter more challenging to detect in our experimental design).

Due to how this experiment was administered, we were not able to determine when boundaries were placed, but future investigations could examine the specific situational features (e.g., social information) that drive segmentation at the individual-level and how this links to specific aspects of experiential diversity. Prior work has shown that lonely individuals show enhanced vigilance to negative social information^46^, suggesting that reduced social experiential diversity may increase sensitivity to event boundaries. However, it is important to note that loneliness did not significantly correlate with event segmentation in the current study, and – notably – the relationship between experiential diversity and segmentation ability held when controlling for loneliness in our analyses. This is important as it implies that reductions in social contact alone may be sufficient to impose real-world psychological costs, independent of subjective feelings of loneliness.

Experiential diversity may also influence event segmentation by altering the functional integrity of a core brain network linked to event cognition^3,30^. Studies in rodents have reported significant declines in spatial memory when compared to those housed in groups, and this is associated with a host of structural changes in the hippocampus and prefrontal cortex (including demyelination and neuroinflammation)^47,48^. Further, changes in these markers may also be mitigated by later exposure to enriched spatial environments^49^, which may even reverse the effects of isolation on hippocampal plasticity^50^. Human studies similarly support the influence of experiential diversity on this brain network (see e.g., ref. ^51^). For instance, sudden and prolonged isolation (i.e., in polar expeditioners) is linked to lower hippocampal volumes relative to controls^52^. Additionally, engaging in complex exploratory behaviour (e.g., in the local neighbourhood) has been linked to positive affect, and increased hippocampal-striatal connectivity^16^. Together, this cross-species evidence underlines the close relationship between immediate social and/or spatial experience and the brain’s event processing system^28,56–58^.

It is important to note that this study was conducted as COVID-19 restrictions were being gradually lifted in the United Kingdom, meaning that many children and adults had undergone a prolonged and acute phase of isolation, with restrictions on social interaction, work, education, and travel. As such, individuals who scored low on our experiential diversity measure (covering 30 days prior to test) may well have undergone an even more sustained period of low experiential diversity. Given the well-established relationship between stress and cognition^59^, the effects in the current study may be stronger than would be typically observed in this population due to prolonged exposure to environmental stressors. Animal studies have shown that stress significantly influences global remapping in hippocampal place cells – specifically, an inability to efficiently shift spatial representations across different contexts^60^. It is possible, therefore, that individuals with low experiential diversity, demonstrated stress-related reductions representational flexibility when faced with changing spatial and/or social environmental features, as captured by fewer perceived event boundaries (though, critically, our results held when accounting for anxiety).

Overall, this study demonstrates a novel link between experiential diversity and event segmentation granularity, offering new insights into how real-world variation in social and environmental connectedness may shape cognition. Understanding these relationships has significant implications for interventions aimed at enhancing cognitive health through increased experiential diversity. Future work should focus on better understanding how experiential diversity shapes encoding of different forms of event-related information (e.g., social vs. non-social information) across the lifespan, and how this relates to ‘downstream’ aspects of cognition, including episodic memory and navigation.

## Acknowledgements

This work was supported by the Biotechnology and Biological Sciences Research Council (BBSRC) [BB/V010549/1] and internal funding from the Department of Psychology, Royal Holloway, University of London. For the purpose of open access, the author has applied a Creative Commons Attribution (CC BY) licence to any Author Accepted Manuscript version arising. We would also like to thank Saloni Krishnan, Nura Sidarus, and Andrew Lawrence for their valuable discussion.

## CRediT author statement

**Carl J. Hodgetts:** Conceptualization, Investigation, Formal Analysis, Data Curation, Project Administration, Visualization, Writing – Original Draft, Writing – Reviewing and Editing. **Samuel C. Berry:** Conceptualisation, Writing – Reviewing and Editing. **Mark Postans:** Conceptualization, Investigation, Formal Analysis, Writing – Reviewing and Editing; **Angharad N. Williams:** Conceptualization, Investigation, Formal Analysis, Writing – Reviewing and Editing.

