## Supplementary material for "Individual differences in experiential diversity shape event segmentation granularity"

### Supplementary information for “Individual differences in experiential diversity shape event segmentation granularity”

This file contains:

- Appendix A - Social experiences questionnaire
- Appendix B - Spatial experiences questionnaire

### Appendix A

#### Social experiences questionnaire

Considering your social interactions over the last month (i.e., 30 days), please indicate whether the following statements apply to you or not.

In the last month....

| Question | Option 1 | Option 2 |
|---|---|---|
| 1. I have lived alone | No | Yes |
| 2. I have had regular (i.e., weekly) face-to-face contact with close family or friends | No | Yes |
| 3. I have had regular (i.e., weekly) voice or video call contact with close family or friends? | No | Yes |
| 4. I have regularly attended a workplace, community group, sports club, and/or volunteering scheme with other individuals | No | Yes |
| 5. I have regularly used messaging apps and social media to stay in touch with close family or friends | No | Yes |
| 6. I have had regular contact with other individuals in my local community who are not primarily my friends of family (e.g., carer, delivery person, shopkeeper, neighbour, plumber, etc) | No | Yes |

##### Scoring

Each item is scored 1 point for the ‘Yes’ response, except for Item 1, which is reverse coded (1 point = ‘No’). Scores are between 0 and 6.

#### Appendix B

##### Spatial experiences questionnaire

| Question | Response options | | | | |
|---|---|---|---|---|---|
| On a typical day within the last month (30 days), how many rooms did you spend your time in? | 1 | 2-4 | 5-7 | 8+ | |
| Do you have access to a private outdoor space? | No | Yes | | | |
| On a typical week in last month, how often did you leave your home (e.g., for exercise, shopping, errands)? | 0 | 1 | 2-4 | 5-7 | 7+ |
| Within the last month (30 days), how often have you visited recreational green space (e.g., woodland, parkland, forest)? | Never | At least once a month | At least once a fortnight | At least once a week | Every day |
| In the last month (30 days), how often have you taken a leisurely wander or stroll without any clear aim or goal? | Never | At least once a month | At least once a fortnight | At least once a week | Every day |
| **Score allocated** | 0 | 1 | 2 | 3 | 4 |
